# Combinatorial native MS and LC-MS/MS approach reveals high intrinsic phosphorylation of human Tau but minimal levels of other key modifications

**DOI:** 10.1101/2020.09.03.281907

**Authors:** Friedel Drepper, Jacek Biernat, Senthillvelrajan Kaniyappan, Helmut E. Meyer, Eva Maria Mandelkow, Bettina Warscheid, Eckhard Mandelkow

**Affiliations:** Biochemistry and Functional Proteomics, Institute of Biology II, University of Freiburg, Schänzlestr. 1, 79104 Freiburg, Germany; Signalling Research Centres BIOSS and CIBSS, University of Freiburg; DZNE (German Center for Neurodegenerative Diseases), Venusberg-Campus 1, Building 99, 53127 Bonn, Germany; Department of Neurodegenerative Diseases and Geriatric Psychiatry, University of Bonn, Venusberg-Campus 1, Building 82, 53127 Bonn, Germany; Medical Proteome -Center, Ruhr-University Bochum, Universitätsstraße 150, 44801 Bochum, Germany; Leibniz-Institute für Analytical Sciences (ISAS), Biomedical Research Dortmund, Germany; CAESAR Research Center, Ludwig-Erhard-Allee 2, 53127 Bonn, Germany

**Keywords:** Tau protein, native mass spectrometry, LC-MS, phosphorylation, Alzheimer disease

## Abstract

Abnormal changes in the neuronal microtubule-associated protein Tau, such as hyperphosphorylation and aggregation, are considered hallmarks of cognitive deficits in Alzheimer disease. Hyperphosphorylation is thought to take place before aggregation, and therefore it is often assumed that phosphorylation predisposes Tau towards aggregation. However, the nature and extent of phosphorylation has remained ill-defined. Tau protein contains up to 85 potential phosphorylation sites (80 Ser/Thr, and 5 Tyr P-sites), many of which can be phosphorylated by various kinases because the unfolded structure of Tau makes them accessible. However, limitations in methods (e.g. in mass spectrometry of phosphorylated peptides, or antibodies against phospho-epitopes) have led to conflicting results regarding the overall degree of phosphorylation of Tau in cells. Here we present results from a new approach, that is based on native mass spectrometry analysis of intact Tau expressed in a eukaryotic cell system (Sf9) which reveals Tau in different phosphorylation states. The extent of phosphorylation is remarkably heterogeneous with up to ∼20 phosphates (P_i_) per molecule and distributed over 51 sites (including all P-sites published so far and additional 18 P-sites). The medium phosphorylated fraction P_m_ showed overall occupancies centered at 8 P_i_ (± 5 P_i_) with a bell-shaped distribution, the highly phosphorylated fraction P_h_ had 14 P_i_ (± 6 P_i_). The distribution of sites was remarkably asymmetric (with 71% of all P-sites located in the C-terminal half of Tau). All phosphorylation sites were on Ser or Thr residues, but none on Tyr. Other known posttranslational modifications of Tau were near or below our detection limit (e.g. acetylation, ubiquitination). None of the Tau fractions self-assemble readily, arguing that Tau aggregation is not promoted by phosphorylation per se but requires additional factors.

## Introduction

Tau (*MAPT*, Uniprot P10636) is a developmentally regulated protein promoting microtubule-based functions like axonal transport in neurons. The properties of Tau can be altered by various post-translational modifications (PTMs), most conspicuously by phosphorylation (Wang and Mandelkow, 2016). The aggregation of Tau into amyloid-like insoluble filaments is one of the hallmarks of neurodegenerative diseases called tauopathies, including Alzheimer disease (AD). Early observations suggested a high level of phosphorylation of Tau preceding that of aggregation (Grundke-Iqbal et al., 1986, Braak et al., 1994) which was therefore interpreted to promote aggregation and is used as an early marker of neuronal degeneration. Tau-targeted therapies, aiming at reducing Tau levels, Tau distribution or Tau modifications have emerged as potential strategies for treating tauopathy in patients (DeVos et al., 2017, Yanamandra et al., 2013, Sigurdsson, 2018). Likewise, quantitative diagnostics including the analysis of soluble Tau from cerebrospinal fluid is gaining importance (Barthelemy et al., 2019). These examples underscore the need for improved quantitative diagnostics.

Initial quantitative assessments of Tau-bound phosphate yielded ∼8 P_i_ per Tau molecule in AD Tau (Ksiezak-Reding et al., 1992, Kopke et al., 1993). This was compared with fetal Tau at ∼7 P_i_ per Tau (Kenessey and Yen, 1993) and adult human cytosolic Tau at ∼2 P_i_ (Ksiezak-Reding et al., 1992, Kopke et al., 1993), interpreted to mean that the extent of Tau phosphorylation is abnormally high in AD. In parallel, phosphorylation dependent antibodies and phospho-peptide mapping approaches were developed to identify phospho-epitopes on Tau (e.g.(Augustinack et al., 2002); for a current list see Alzforum.org). However, these antibody-based methods were not well suited for the determination of the occupancy of the P-sites and the overall state of phosphorylation.

The advent of Edman degradation and mass spectrometry (MS) resulted in increased sensitivity and specificity of detection of PTMs on Tau protein. In particular the combination of high performance liquid chromatography with tandem MS (HPLC-MS/MS) resulted in enhanced coverage of the Tau sequence with newly identified P-sites (review,(Hanger et al., 2009, Hanger, 2020)).

However, to address the issue of hyperphosphorylation versus aggregation, an experimental system was needed which enabled high phosphorylation of defined Tau isoforms. This was achieved with the expression of full-length human Tau (2N4R) in transfected Sf9 cells which yields high protein levels (up to 230 µM in cells) and high diversity of P-sites, as judged by antibody reactivity and MALDI-TOF MS analysis (Tepper et al., 2014). Depending on cell treatment (without or with phosphatase inhibitor okadaic acid), the overall occupancy was estimated around 12 P_i_ or 20 P_i_. Hence the fractions were termed P12 or P20, renamed in this paper to P_m_ and P_h_ to indicate medium versus high phosphorylation state, and in contrast to unphosphorylated P_o_ Tau expressed in *E*.*coli* **(Fig. 1A-C)**. None of the phosphorylated Tau proteins showed an enhanced tendency for aggregation (Tepper et al., 2014).

**Fig. 1:**
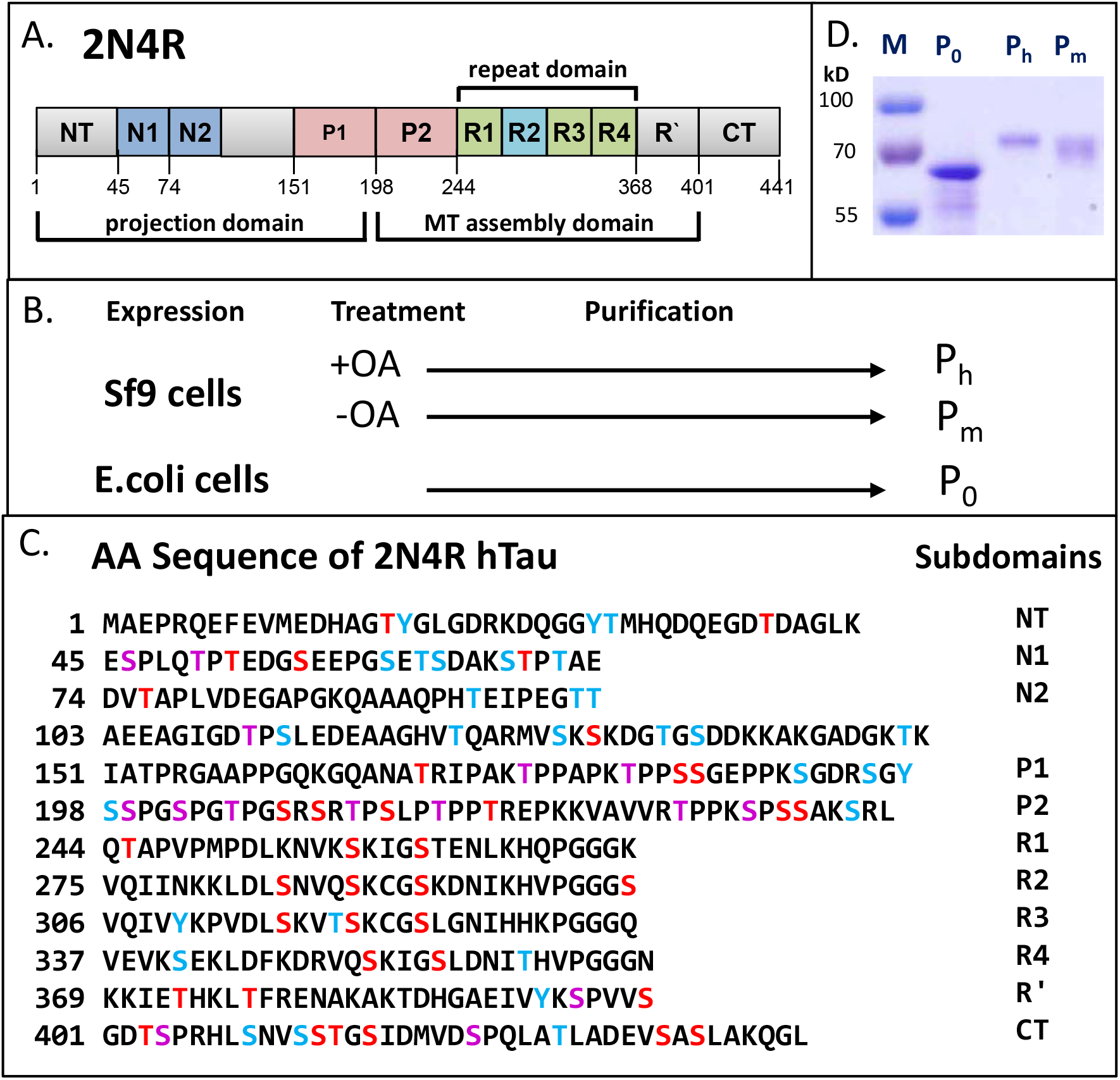
Phosphorylation of hTau-2N4R expressed in Sf9 cells and E.coli Overview of Tau domains, sequence, preparations. Overview: Phosphorylation of hTau-2N4R expressed in Sf9 cells and *E. coli*. ***(A)*** Diagram of domains of hTau40 (2N4R or Tau-F, Uniprot Id P10636-8), the largest human isoform of CNS Tau consisting of 441 residues with the three alternatively spliced inserts N1, N2 and R2. The N-terminal half represents the projection domain, and the C-terminal half contains the four pseudo-repeats (R1-R4) which - in combination with their flanking domains P2 and R’ - represent the microtubule assembly domain. ***(B)*** Schematic representation of production of unphosphorylated Tau (P_0_) in *E. coli* (prokaryote) and hyperphosphorylated Tau (P_m_ and P_h_) Tau in Sf9 cells (eukaryote). Okadaic acid (OA) treatment increases the phosphorylation of Tau which yields P_h_ -Tau. ***(C)*** Amino acid sequence of Tau (2N4R, 441 residues) showing phosphorylation sites identified in this study (*red and purple letters*; *purple=Pro directed)*) and further potential phosphorylation sites (not detected so far, *blue letters*) (total of 85 sites, 45 Ser, 35 Thr, 5 Tyr). Note that only a minority of phosphorylation sites of 37% was detected in the N-terminal part of the sequence (up to residue ∼150, compared to 71% in the C-terminal part of the protein. Note also that none of the five Tyr residues was phosphorylated. ***(D)*** SDS-PAGE analysis stained with Coomassie blue showing P_0_ Tau purified from *E. coli*, P_m_ and P_h_ Tau purified from Sf9 cells. Note the upward shift in Mr value with increasing phosphorylation, from 55 kDa (P_0_-Tau) to 68 kDa for P_h_-Tau (compared with the theoretical MW values of ∼45850 Da and ∼46863 Da). This shift is characteristic for AD-Tau.

Nonetheless, the same fractions were studied by a novel MS-based assay, FLEXITau (Mair et al., 2016), designed to determine the locations and occupancies of all P-sites quantitatively by comparison with isotopically labeled peptide forms. This procedure yielded overall occupancies of only 7 and 8 P_i_ per Tau molecule in P_m_-Tau and P_h_-Tau, respectively, with a broad spread (±5 around the mean), and up to 23 observed P-sites. Other MS approaches, designed to identify P-sites in Tau from AD brains and CSF, revealed ∼30 P-sites but not the overall occupancies (Sato et al., 2018).

In the present work, we employed native MS to determine the extent of Tau phosphorylation in cells and to clarify the variations between different methods. MS of intact proteins, such as top-down MS (Donnelly et al., 2019) and native MS (Boeri Erba and Petosa, 2015, Hernandez and Robinson, 2007), allow quantification of different proteoforms without requiring prior proteolytic cleavage into peptide fragments or comparison with reference standards (Schaffer et al., 2019). The procedure revealed that full-length Tau in the P_m_ fraction contained 8± 5 P_i_, and in the P_h_ fraction 14± 6 P_i_. Subsequent analysis of phosphorylated peptides revealed up to 51 P-sites in Tau, with variable site occupancies. Finally, the method sensitively detected a low level of different cellular proteins associated with Tau.

## Results

### Analysis of intact full-length phospho-Tau by native MS

Analysis of tryptic peptides by HPLC-MS/MS reveals sites of protein phosphorylation, but does not directly monitor the extent of phosphorylation per molecule (i.e. overall occupancy) because the detection efficiency varies between phosphorylated and non-phosphorylated peptides. This problem is circumvented by native MS of full-length Tau (Boeri Erba and Petosa, 2015). We compared the phosphorylation status of two differently phosphorylated Tau fractions expressed in eukaryotic Sf9 cells with unphosphorylated Tau from *E*.*coli* bacteria **(Fig. 1B). Fig. 2** shows native MS spectra of unphosphorylated control Tau (termed P_o_, expressed in *E. coli*, bottom), Tau from the fraction P_m_-Tau (medium level phosphorylation, expressed in Sf9 cells, middle) and highly phosphorylated fraction P_h_-Tau (in presence of phosphatase inhibitor okadaic acid, top). For P_o_-Tau the main signals were recorded between *m/z* 2500 and 3500. By charge state deconvolution a series of signals corresponding to charges +16 to +13 can be related to a molecular weight of 45724 Da (**Fig. 2**, open red circles; see also **Fig. 3**). Native MS spectra of P_m_-Tau and P_h_-Tau show several matching signals in the range below *m/z* 2750 and above *m/z* 3500 (**Fig. 2**, middle and top spectrum). In between, broad peaks are visible for both P_m_-Tau and P_h_-Tau which can be assigned to a series of charge states +16 to +14 (red squares for P_m_-Tau and filled red circles for P_h_-Tau). Both series are shifted toward higher *m/z* compared to the corresponding charge states of the control Tau P_o_. Further peak series in P_o_-Tau were attributed to a trimeric complex (grey circles, bottom trace) of the *E. coli* periplasmic chaperone protein Skp (molecular mass of full-length Skp 17688 Da, reduced to 15691 Da after cleavage of 20 residue signal peptide (see Table 3)), and to an unknown protein of 59642±5 Da (grey squares, top and middle traces) copurified with P_m_-Tau and P_h_-Tau from Sf9 cells.

**Table 1:**
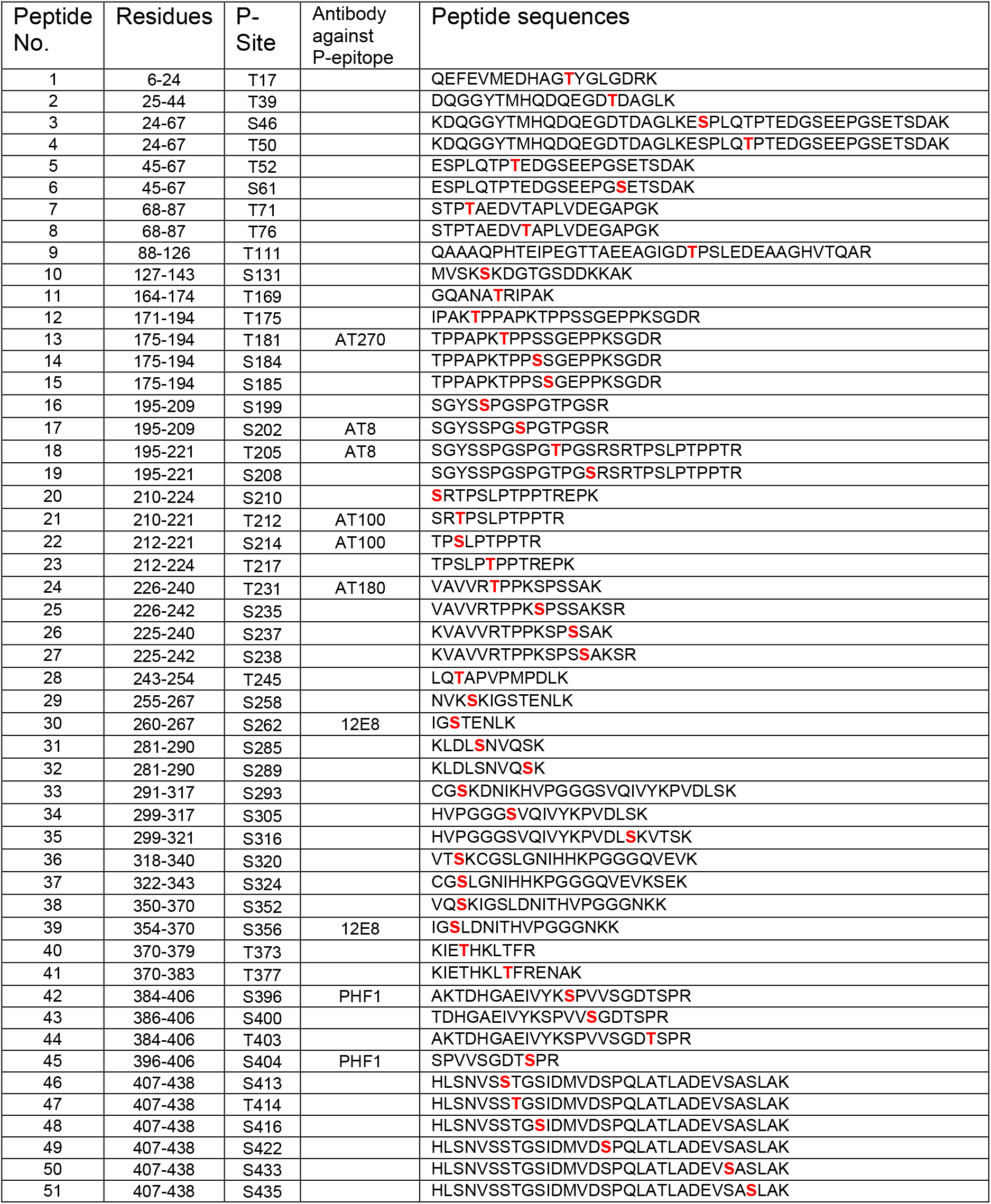
Phosphorylation sites on Tau protein (fractions P_m_-Tau and P_h_-Tau) identified by LC-MS/MS. Note that phosphorylation is observed only on Ser or Thr, not on Tyr.

**Tab. 2:**
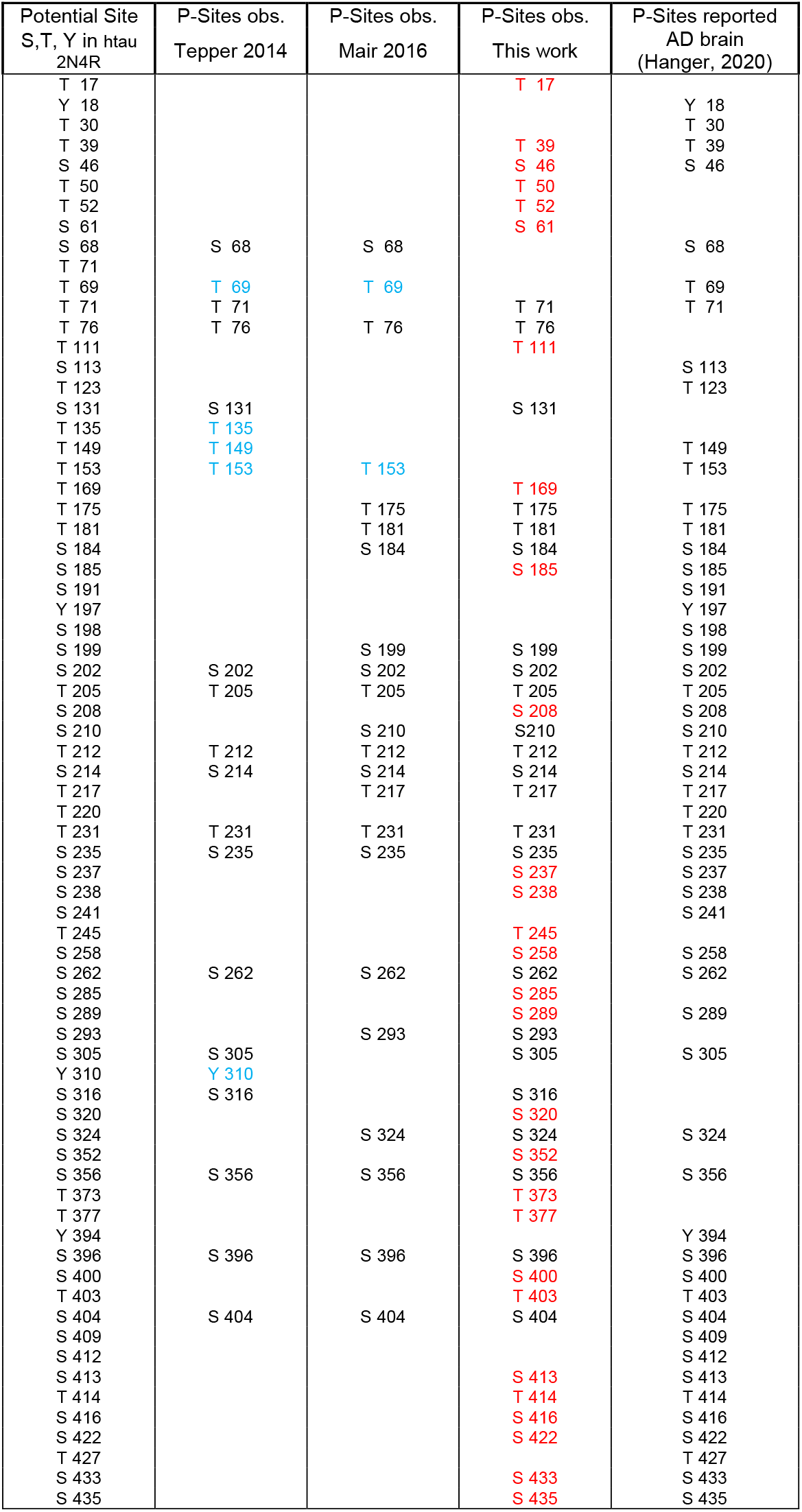
P-sites in human Tau (2N4R) expressed in Sf9 cells by three independent studies: Tepper et al. (2014) [MALDI-TOF MS], Mair et al. (2016) [FLEXITAU] & current work [native MS and LC-MS/MS]. Col 1= potential P-sites in full-length Tau (residues S, T, Y), col 2-4 = observed P-sites in Tau from Sf9 cells, col. 5= P-sites in Tau from human AD brain (Hanger 2020). Red = additional sites observed in current work, blue = sites observed in previous but not in present study.

**Table 3:**
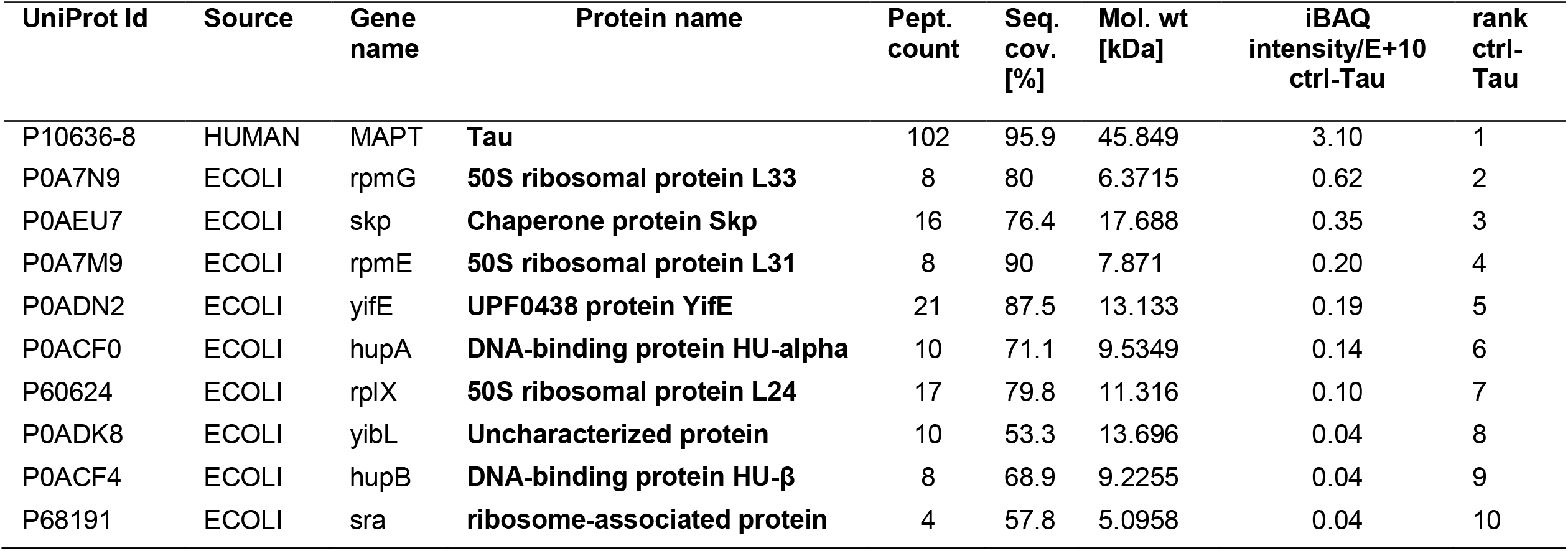
Ten most abundant Tau-associated proteins identified by LC-MS/MS analysis in samples of purified P_o_-Tau expressed in *E*.*coli*. Protein samples were digested with trypsin followed by LC-MS/MS. Data were analyzed using MaxQuant software and Uniprot organism specific databases for *H. sapiens, E*.*coli* and *Spodoptera frugiperda* (SPOFR). iBAQ = intensity-based absolute quantification correcting for differences in the number of predicted peptides per protein. Only the 10 most abundant proteins per sample are listed. Average of two biological replicates. Note the frequency of RNA- or DNA-binding proteins copurified with Tau.

**Table 4A:**
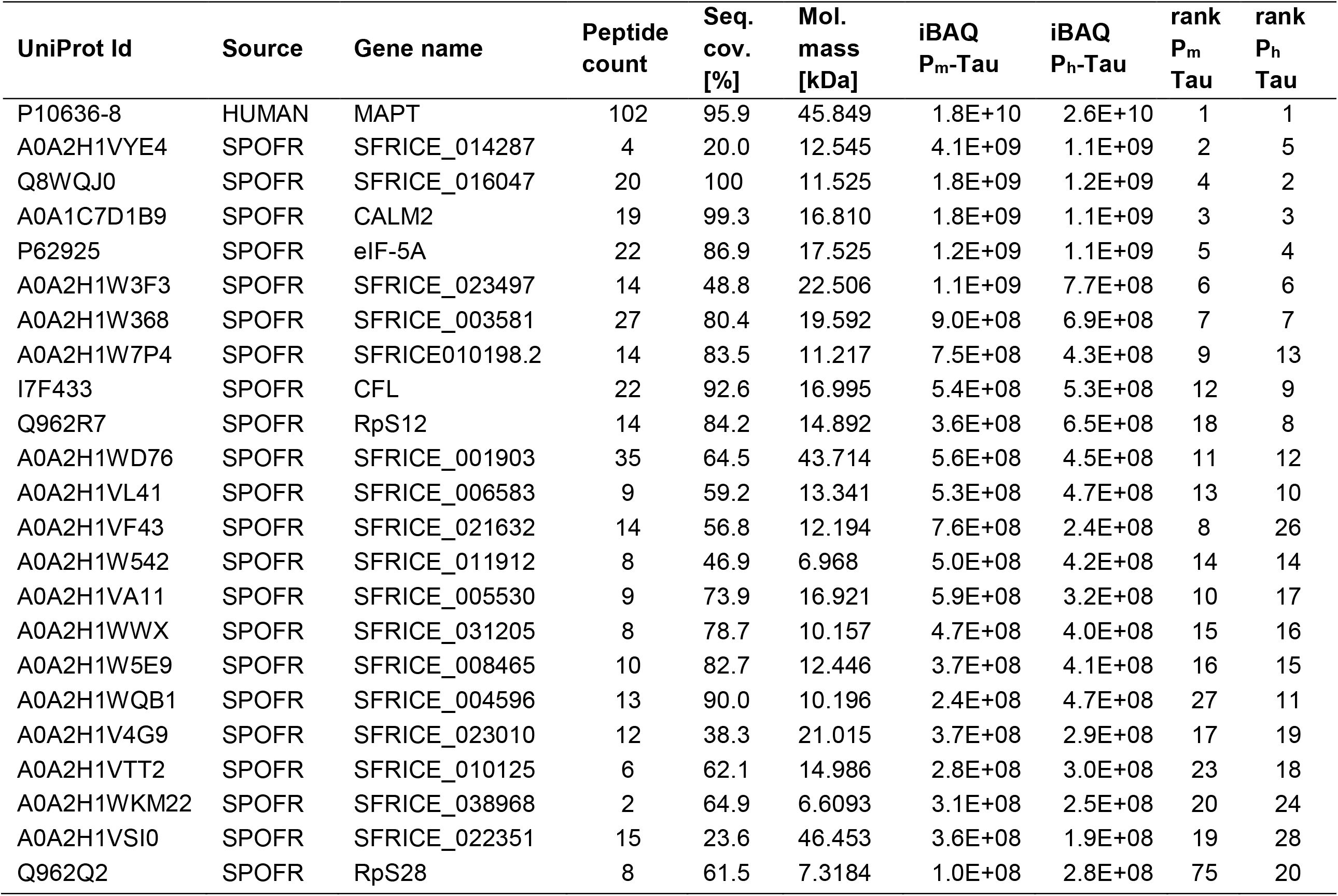
20 most abundant proteins co-purifying with Tau identified by LC-MS/MS in fractions of P_m_-Tau and P_h_-Tau from Sf9 cells. For protein functions see Tab. 4B.

**Table 4B:**
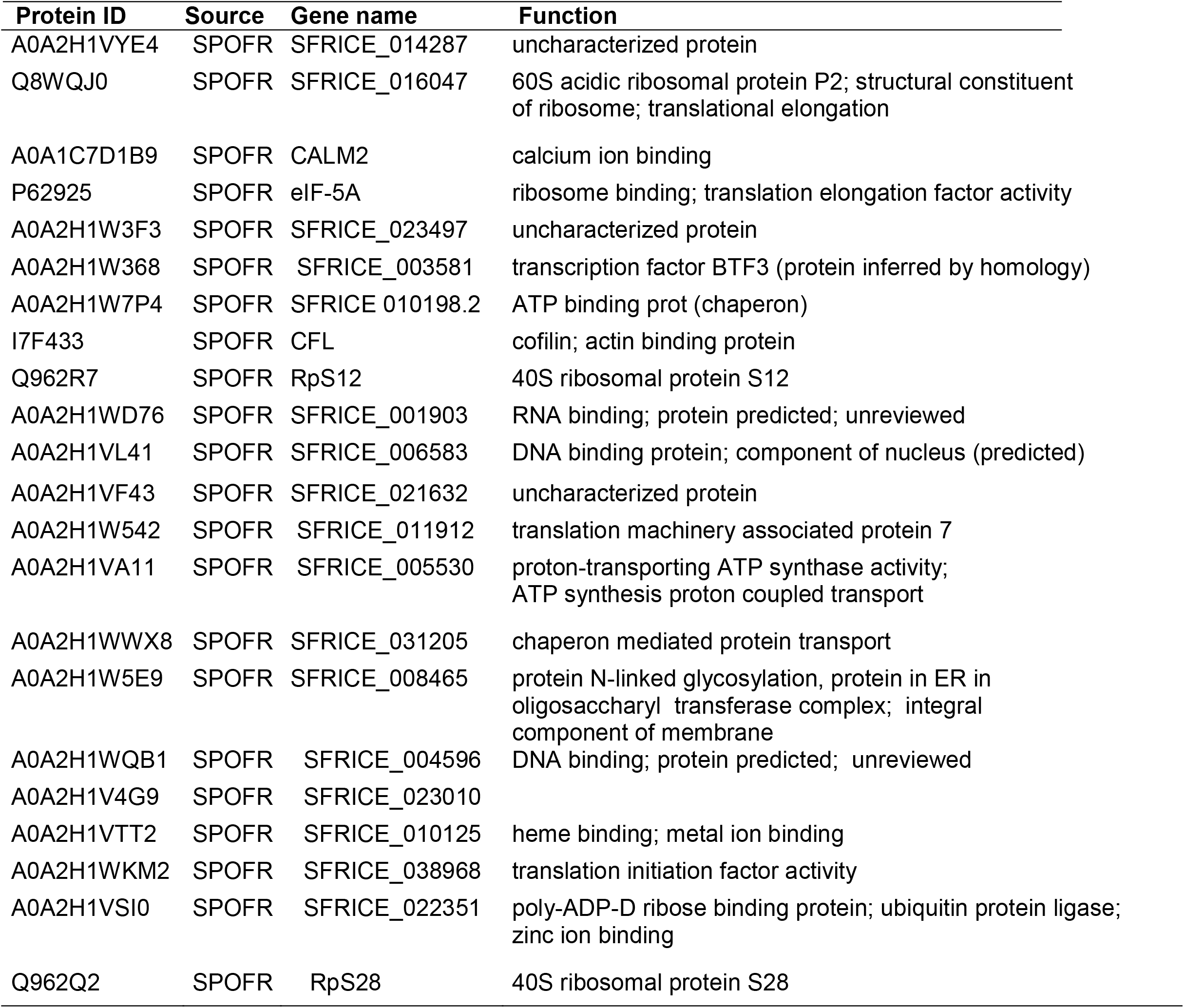
Known functions of proteins co-purified with Tau.

**Fig. 2:**
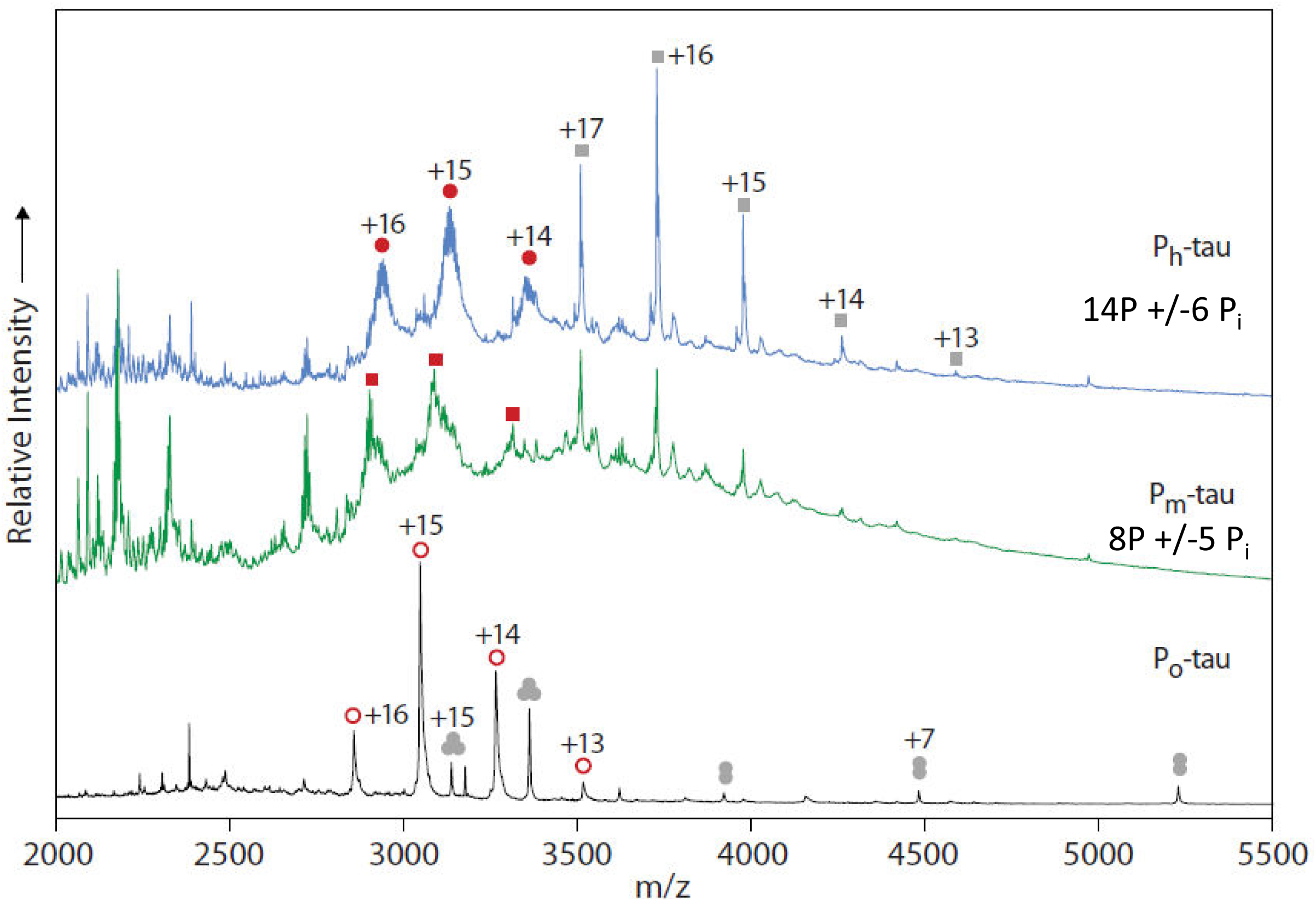
Analysis of hTau-2N4R expressed in Sf9 cells by native MS: charge state assignment. Analysis of hTau-2N4R expressed in Sf9 cells by native MS. 20 µM of purified proteins in 200 mM ammonium acetate, pH 7.6, were analyzed by nanoflow electrospray ionization quadrupole time-of-flight mass spectrometry. Representative mass spectra for the hyperphosphorylated states P_h_-Tau (top) and P_m_-Tau (middle) as well as unphosporylated P_0_-Tau (bottom) are shown. **Top**, main signals were assigned to charge state series of P_h_-Tau (MW of 46883 Da; filled red circles) and a copurified component with a molecular weight of 59642 Da (grey squares). **Middle**, spectrum of P_m_-Tau displaying peaks assigned to a series of equivalent charge states, but shifted towards lower *m/z* compared to those for P_h_-Tau (red squares). Further signals match closely with those in the spectrum of P_h_-Tau. **Bottom**, spectrum of control Tau P_0_ consisting of a series of charge states indicating a molecular weight of 45,724 Da (open red circles). A further series was assigned to a MW of 47,085 Da. It was attributed to a trimeric species (grey circles), as it decomposes upon increasing collisional activation into highly charged (centered at +9, below *m/z* 2,000) monomeric and charge-stripped dimeric species (charges +6 to +8). Its monomeric mass of 15,691 Da matches the theoretical mass of *the E. coli* chaperone protein *skp* (see **Table 3**), taking into account removal of its N-terminal 20 amino acids signal peptide.

**Fig. 3:**
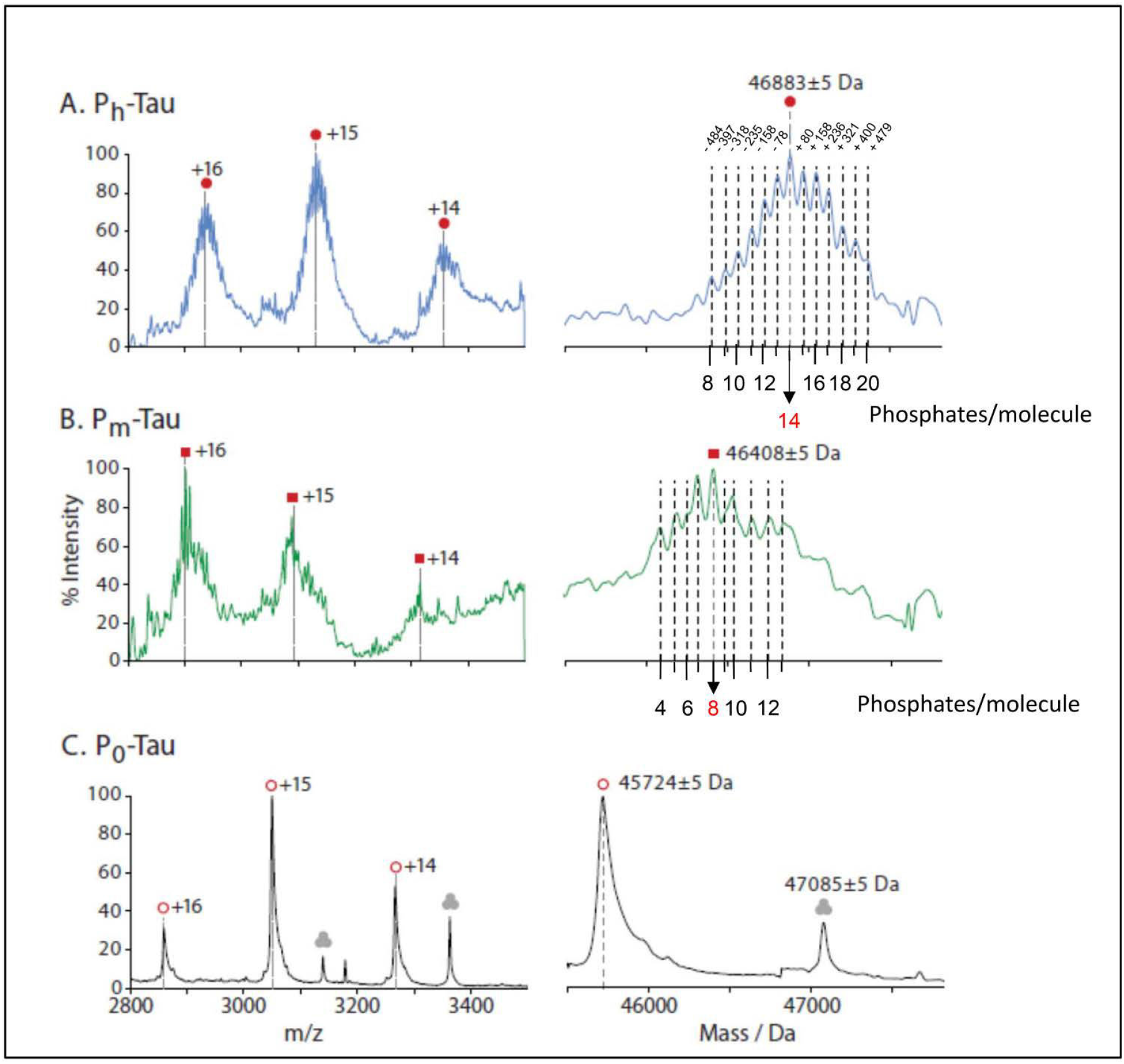
Analysis of hTau-2N4R expressed in Sf9 cells by native MS: charge state-deconvoluted spectra. Phosphorylation of hTau-2N4R expressed in Sf9 cells analyzed by native MS. **Left**, signals in the range between 2800 and 3500 *m/z* attributed to Tau proteins. **Right**, corresponding charge state-deconvoluted spectra. The mass spectrum of P_h_-Tau (top) displays a center mass of 46,883 Da (filed red circles) and additionally six peaks on both sides equally spaced by ∼80 Da. P_m_-Tau (middle) displays peaks of similar width shifted towards lower *m/z* compared to those for P_h_-Tau, which can be assigned to the equivalent charge states and indicate a molecular weight of approximately 46,408 Da (red squares). The spectrum of control Tau P_0_ consists of a series of charge states resulting in a comparatively sharp peak at a mass of 45,724 Da (45,850 -131Da +/- 5 of missing Met-1; open red circles). A further series was assigned to a MW of 47,085 Da and was attributed to a trimeric species (grey circles), as it decomposes upon increasing collisional activation into highly charged (centered at +9, below *m/z* 2,000) monomeric and charge-stripped dimeric species (charges +6 to +8). Its monomeric mass of 15,691 Da matches the theoretical mass of *the E. coli* chaperone protein *skp* (see **Table 3**), taking into account removal of its N-terminal 20 amino acids signal peptide.

Native MS spectra in the range of *m/z* 2800 to 3500 (**Fig. 3**, left) were analyzed by charge state deconvolution to generate zero-charge mass spectra revealing further differences between these Tau protein forms (**Fig. 3**, right). For P_o_-Tau from *E*.*coli* (bottom), a sharp peak indicates a molecular weight of 45724 Da which is 126 Da lower than the MW of unmodified full-length Tau (2N4R, calculated at 45850 Da). Taking into account the mass accuracy of ±5 Da this can be explained by a cleavage of the N-terminal methionine (131 Da; Fig. 3). For P_h_-Tau from Sf9 cells (**Fig. 3**, top), a broader peak in the zero-charge mass spectrum displays a center mass of 46883 Da and up to six additional peaks on both sides equally spaced by approximately 80 Da. The center mass would correspond to an N-terminally acetylated htau40 carrying 14±6 P_i_ groups.

The spectrum of P_m_-Tau was less well resolved and possibly disturbed by a feature at *m/z* 2750, which is present in the P_h_-Tau spectrum at much lower intensity. A charge state deconvolution results in a center mass of 46408 Da ± 400 Da and, indicating 8±5 P_i_ (Fig. 3, middle row). Of note, although the native MS spectrum of P_m_-Tau was less well defined, all three charge states of +16 to +14 are clearly shifted towards lower *m/z* values compared to the P_h_-Tau spectrum, but remained at higher *m/z* values compared to the corresponding charge states in the control spectrum of P_0_ -Tau (see **Fig. 2** and **Fig. 3**).

### Identification of peptide sequences and posttranslational modifications by LC-MS/MS

#### (a) Phosphorylation

For protein identification and detection of PTMs the protein samples were subjected to reduction of cysteine residues, alkylation of free thiol groups and digestion with trypsin followed by LC-MS/MS analysis. The identified tryptic peptides covered 95% of the Tau sequence and allowed us to detect sites of posttranslational modifications of the Tau proteins. We therefore analyzed our LC-MS/MS data for the following amino acid modifications: acetylation of the protein N-terminus, phosphorylation of serine, threonine or tyrosine, acetylation of lysine, and ubiquitination of lysine. For purified P_m_-Tau and P_h_-Tau, only N-terminally acetylated forms of peptides containing the protein N-terminus (without Met1) were detected. In contrast, for control P_0_-Tau these two peptides were identified in their non-modified, non-acetylated N-terminal peptide form. We therefore deduced from the protein sequence (residues 2-441) average masses of 45718.2 Da for the non-N-terminally acetylated *E*.*coli* Tau (P_0_) and 45760.2 Da for the N-terminally acetylated form of htau40 (P_m_ and P_h_) in the absence of further modifications.

The total number of P-sites on serine and threonine residues on Tau identified at a localization probability higher than 0.75 was 51 sites (32 Ser, 19 Thr). No phospho-tyrosines were detected (for P-sites see Table 1; for procedures to derive site probabilities see Methods). No P-sites were detected for *E. coli* P_0_-Tau, as expected. The P-site analysis of Tau from Sf9 cells confirmed the majority of P-sites found in previous studies (Mair et al., 2016, Tepper et al., 2014), including all prominent Alzheimer specific antibody phospho-epitopes, notably those that require pairs of phosphorylated residues (e.g. antibodies AT-8, PHF-1, AT100). Furthermore, the analysis revealed ten additional P-sites (T373, T377, S400, T403, S413, T414, S416, S422, S433, S435, see Table 1) in the C-terminal domain of Tau, which have been reported for Tau from human AD brain (reviews (Hanger et al., 2009, Hanger, 2020)) but were absent in our previous studies. Interestingly six P-sites (T245, S285, S293, S316, S320, S352, see Table 2) were identified in the repeat domain which are not annotated in the collection of P-sites from AD brains (Hanger et al., 2009, Hanger, 2020). The highly phosphorylated fraction P_h_ contains 14 Pi (± 6 Pi) per Tau molecule indicating a maximal value of 20 Pi (higher values are possible, but not resolved in the spectrum). The heterogenous P-site distribution over the entire length of the Tau chain is generally in line with previous findings (Mair et al., 2016)(and see Discussion).

#### (b) Other posttranslational modifications

In contrast to the many identified P-sites, it is remarkable that other well-documented PTMs of Tau reported for human and mouse brain are close or below the detection threshold in Tau from Sf9 cells. This includes lysine acetylation and lysine ubiquitination, both of which have been linked to neurodegeneration in AD (Min et al., 2010, Cohen et al., 2011, Arakhamia et al., 2020) as well as O-GlcNAc modification of serine and threonine, which was reported to prevent phosphorylation and aggregation of Tau (Hart et al., 2011, Yuzwa et al., 2012). Of note, in contrast to the high number of detected P-sites and their high abundancies, other PTMs, if significant at all, were observed only sporadically and in low intensity. Further validation and characterization of such low abundant additional modifications would need specialized approaches and is beyond the scope of the present study.

#### (c) Tau from neuronal origin

In addition to Tau expressed in cell models we studied the phosphorylation status of Tau isolated from brains of transgenic mice, using LC-MS/MS and native MS methods. The LC-MS/MS analysis was performed on neuronal Tau from transgenic mice expressing the mutation Tau^A152T^ (a risk factor for PSP, (Coppola et al., 2012)) purified by immunoprecipitation using the Tau^A152T^ specific antibody 1C5. The analysis showed that human Tau^A152T^ expressed in mice is highly phosphorylated at five sites (T181, T231, S199, S202, S396; P-site probability > 0.75). These sites are consistent with the subset of P-sites in Sf9 cells with the highest occupancy (>50%) (Mair et al., 2016) and with prominent P-sites of Tau in AD brain. However, native-MS did not reveal a specific spectrum of Tau from neuronal origin, mainly because of the inherent heterogeneity of neuronal Tau proteins, low concentration and contamination by other proteins. This problem will require further development.

### Proteins co-purifying with Tau identified by LC-MS/MS

The high sensitivity of LC-MS/MS revealed the presence tryptic peptides originating from traces of other proteins accompanying Tau through the purification procedure. The recombinant Tau proteins (P_0_ from *E*.*coli*, P_m_ and P_h_-Tau from Sf9 cells) were purified by making use of the heat stability of Tau protein whereby nearly all other proteins are denatured and precipitated whereas Tau remains in solution because of its hydrophilic nature. The detected small amounts of cellular Tau-binding proteins also passed through the boiling step and were still present in gel filtration column fractions containing the separated Tau proteins. One possibility to explain the presence of these proteins is their high affinity for Tau protein. Thus, the experiments yielded a list of high-affinity Tau-interacting proteins which might become interesting for analyzing possible Tau interactions in other cell types (see **Table 3** for P_0_-Tau from *E. coli* and **Table 4** for P_m_-Tau and P_h_-Tau from Sf9 cells). A substantial part of identified proteins are known for their roles in interacting with RNA or DNA, and with chaperones (see Discussion).

## Discussion

### Significance of Tau phosphorylation in AD research

Tau is a neuronal protein whose best-known role is to stabilize microtubules in axons, thus supporting their role as tracks for axonal transport and as struts of axonal shape (Drubin et al., 1985). Tau has a hydrophilic character, contains many charged residues, is highly soluble and has a mostly unfolded structure; as such it can interact with many cellular components in addition to microtubules (Wang and Mandelkow, 2016). The interactions can be modulated by various PTMs, notably by phosphorylation at Ser, Thr, or Tyr residues (up to 85 potential sites). It has been reported that in physiological conditions human fetal Tau is highly phosphorylated (∼7-10 Pi per Tau molecule), compared with human adult Tau (∼2 Pi), and high again for human AD patients (7-10 Pi) (Kenessey and Yen, 1993, Kopke et al., 1993, Ksiezak-Reding et al., 1992). However, these values need to be re-interpreted in the light of later discoveries (see below). In hibernating animals, Tau becomes highly phosphorylated in a reversible manner, corresponding to a cyclic regression and reappearance of dendritic trees, which indicates that Tau phosphorylation may play a role for neuronal plasticity (Arendt et al., 2003). Many cell and animal models of Tau have been developed to study its functions. In particular, Sf9 cells served as early examples of Tau’s role in the control of process outgrowth via microtubules and actin filaments (Baas et al., 1991, Knops et al., 1991) and the importance of reversible phosphorylation (Biernat et al., 2002). Beyond cell biological issues, the major interest in Tau arises from its property as a hallmark of brain diseases, notably Alzheimer and other tauopathies (DeTure and Dickson, 2019). The finding that Tau becomes aggregated into amyloid filaments in AD and appears hyperphosphorylated triggered an extended search for kinases and phosphatases responsible for Tau’s pathological state [reviews, (Martin et al., 2013, Hoffman et al., 2017)], as well as searches for P-sites on Tau (Hanger, 2020), and treatments based on these results (Schneider and Mandelkow, 2008). The aim was to identify P-sites as early indicators of pathology in brain tissue, CSF, or in blood (Barthelemy et al., 2020, Bateman et al., 2020), and as indicators of vulnerable brain circuits affected during Braak stages (Arnsten et al., 2019). In spite of this progress, the relationship between Tau’s phosphorylation and aggregation remains enigmatic in view of its exceptional solubility.

## Methods development

In the past, the main tools for identifying phoshorylation sites of proteins were based on tryptic cleavage, combined either with thin-layer chromatography of ^32^P-labeled phosphopeptides, peptide sequencing (Boyle et al., 1991), or mass spectrometry of peptides (e.g. MALDI-TOF MS) (Hasegawa et al., 1992). For the case of Tau, these methods yielded information on P-sites involved in AD (e.g. (Gustke et al., 1992, Illenberger et al., 1998, Tepper et al., 2014)). A limitation common to these methods was that the extent of phosphorylation of peptides or the whole protein remained uncertain. For peptides, this was partially overcome by using isotope-labeled standards (e.g. (Mair et al., 2016, Barthelemy et al., 2016)). However, the correlation between P-sites and global extent of phosphorylation remained a problem. The early studies of global phospho-occupancy, obtained by ashing and color reaction of samples, revealed average global occupancies but carried no information on occupancy of individual P-sites. Yet knowledge of both the degree of phosphorylation and the main P-sites would be important for assessing the cellular functions (e.g. control of microtubule assembly, phase separation in the cytoplasm) and the pathological state of Tau (e.g. somatodendritic missorting, pathological aggregation). This can be achieved by combining native MS with LC-MS/MS, as described here. We will restrict our discussion to some issues that have received attention in the Tau field during recent years.

### Overall extent of phosphorylation

In previous work (Tepper et al., 2014), we had described three major fractions of phospho-Tau, termed P0, P12, and P20, whose degree of phosphorylation was estimated from the peaks observed by MALDI-TOF MS. Fraction P0 corresponded to the unphosphorylated protein expressed in *E*.*coli*. Fraction P12 was derived from Sf9 cells after expression of full-length Tau (2N4R) whose mass was consistent with additional ∼12 Pi. Fraction P20 was also from Sf9 cells, with phosphatase inhibitor okadaic acid added during preparation, consistent with additional ∼20 Pi.

The same types of fractions were studied by Steen and colleagues (Mair et al., 2016) using the FLEXITau method where quantification was done with reference to isotope-labeled full-length Tau. The fractions were denoted as P-Tau (equivalent to the P12 preparation of (Tepper et al., 2014)), but with a lower overall occupancy of 7 P_i_, and PP-Tau (equivalent to P20 preparation) with an overall occupancy of 8 P_i_. In addition, the method revealed the individual occupancies of 17 sites, including those of the major Tau antibodies (Fig. 3, 4 in (Mair et al., 2016)).

**Fig. 4:**
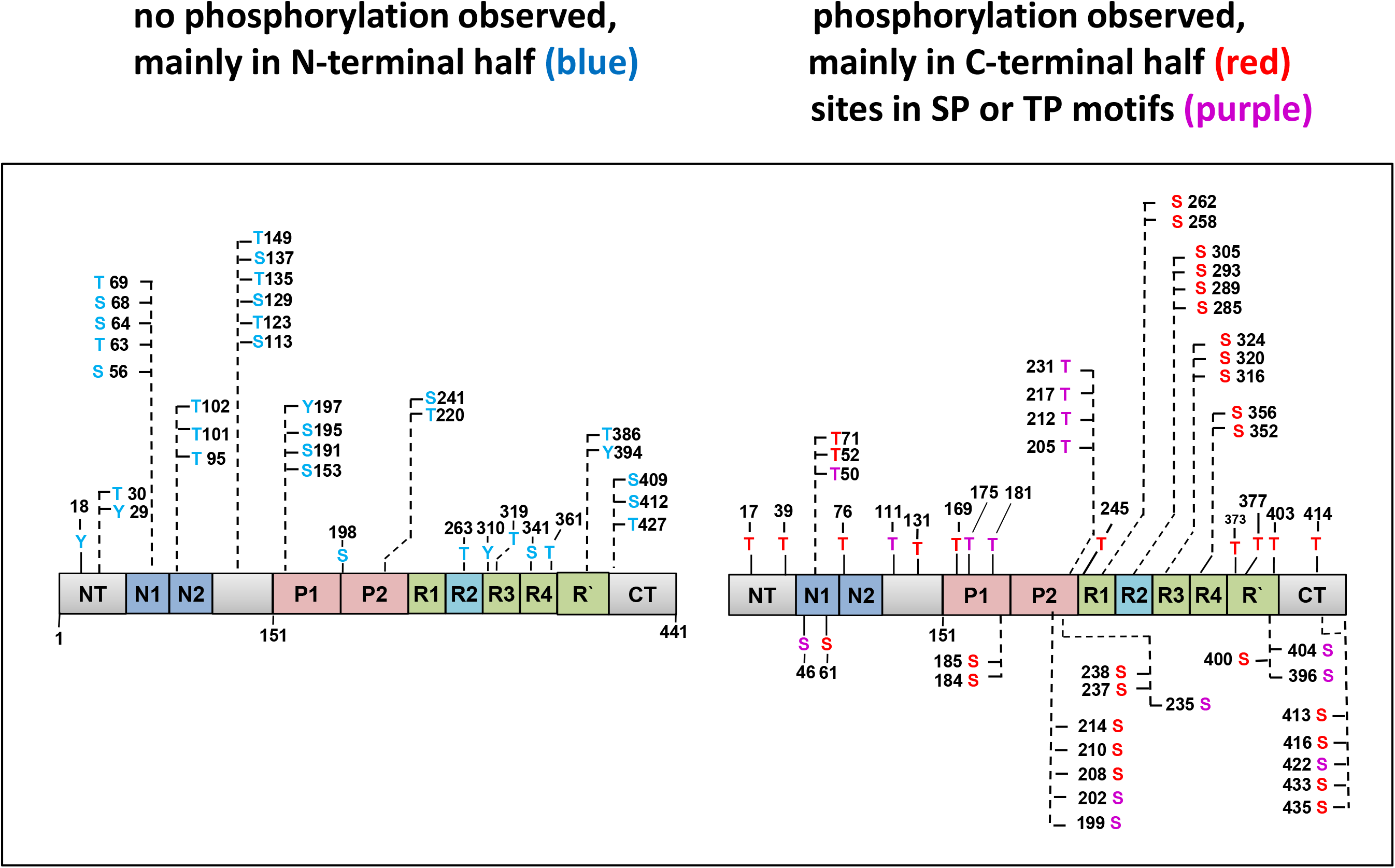
Potential vs observed phosphorylation sites of Tau-2N4R from Sf9 cells. Bar diagram of full-length human Tau protein and distribution of phosphorylation sites. Left, no phosphorylation observed; right, phosphorylation sites observed in Sf9 cells.

The apparent discrepancies between the overall occupancies observed in the earlier MALDI-TOF MS and FLEXITau experiments are now superseded by the results from native MS. The fractions are now denoted as P_0_, P_m_, and P_h_ for zero, medium, and high state of phosphorylation (in order to distinguish the different numerical values). The common control value is P_0_-Tau from *E*.*coli* (Fig. 3C), which remains without modification, apart from N-terminal processing by removal of Met 1. The medium state, reflecting normal kinase/PPase activity in cells (Fig. 3B), is now resolved into a series of peaks centered around a maximum at +8 P_i_ per molecule, with ∼4 sub-maxima resolved on either side (range +4 to +13 P_i_). The high state (Fig. 3A), reflecting normal kinase but inhibited PPase activity, appears as a series of peaks centered around a maximum at +14 P_i_, with ∼6 sub-maxima resolved on either side (range +6 to +20 P_i_).

The main conclusions are: The previous MALDI-TOF MS values agree roughly with the upper limits of the peaks resolved by native MS, the FLEXITau values reflect the lower limits. Of note, a closer inspection of the MALDI-TOF MS results (Tepper et al., 2014) shows that the experimental errors of the reported masses were up to +/-195 Da, corresponding to +/-2.5 P_i_ for individual fractions and thus +/-5 P_i_ for mass differences. On the other hand, the FLEXITau method achieved accurate occupancies of individual P-sites. However, sequence coverage was limited (75%) so that the degree of Tau phosphorylation was underestimated (Table 2, Table S1; note that the newly covered sequences in this study contain 14 additional P-sites).

It is remarkable that Tau molecules in the many P-states are separated by 1 P_i_ unit. For a multi-domain protein (or multi-protein complex) with specific functional P-sites one would expect well-defined stoichiometries of phosphorylation (by specific kinases) whereby assembly or activity are controlled; examples are ribosomes or proteasomes (Guo et al., 2017, Mikulik et al., 2011). By contrast, in case of Tau all domains have a natively unfolded character, with an unusually high fraction of phosphorylatable residues (85/441=19%) accessible to multiple kinases and phosphatases. The degree of phosphorylation therefore appears to depend on a phosphorylation “tone” rather than on specific functional modifications. The balance between kinase and phosphatase activities depends on the experimental conditions of Tau purification which can change rapidly. Thus, during postmortem delay, ATP is depleted (making kinases inactive), and PP2A_cat_ is activated, which makes adult human Tau appear to be in a low state of phosphorylation. Conversely, rapid isolation of brain tissue (e.g. with fetal Tau or brain biopsy) reveals a high state of phosphorylation. In this scheme, the high phosphorylation of AD-Tau (in spite of a long postmortem delay) is explained by Tau aggregation which protects them against PPases. Similarly, lowering the temperature (as in anesthesia) decreases PPase activity, with a lesser effect on kinases, causing the impression of hyperphosphorylation (Planel et al., 2007, Matsuo et al., 1994, Wang et al., 2015). Thus, although there are specific functional or diagnostic P-sites in Tau they probably have only a small effect on the average degree of phosphorylation. By implication, the changes in phosphorylation tone would affect many cellular proteins, not just Tau, so that other proteins might also serve as diagnostic markers for AD neurodegeneration, even when they do not appear in aggregated form.

If multiple phosphorylation of a natively unfolded protein like Tau is largely a statistical property, it follows that each state of phosphorylation represents an ensemble of many Tau species, phosphorylated at different sites, depending on activity and accessibility of kinases/PPases. Certain sites may appear as reliable markers of disease, but this is not necessarily coupled to a specific function (such as aggregation or MT binding). Consistent with this, we find that none of the phosphorylation states led to a pronounced increase in Tau aggregation (Tepper et al., 2014) even though Tau aggregation and phosphorylation seem to be co-regulated during AD progression. However, the extent of phosphorylation will change some bulk properties such as charge, which could turn from net positive to net negative. This would change the interactions with other proteins (e.g. the cytoskeleton via microtubules and actin filaments), RNAs, or membrane surfaces, as well as the folding of Tau from a “paperclip” to other conformations (Jeganathan et al., 2008). In particular, phosphorylation promotes the propensity of Tau to undergo liquid-liquid phase transitions (Wegmann et al., 2018).

### Asymmetric distribution and hotspots of phosphorylation

The sequence of Tau contains several points where phosphorylatable residues are clustered and are phosphorylated, either with high occupancy at a single residue, or with intermediate occupancy at two or three nearby residues. Some of these hotspots of phosphorylation are prominent as epitopes of antibodies raised against AD-Tau. The antibodies may bind to a single site within a cluster, independently of nearby P-sites (e.g. pT181 by antibody AT270, or pS231 by antibody AT180 (Amniai et al., 2011), or require a pair of nearby phosphorylated residues (e.g. S202+T205 (AT8, (Goedert et al., 1995)), S212+S214 (AT100, (Zheng-Fischhofer et al., 1998)), S396+S404 (PHF1, (Otvos et al., 1994)). These clusters are represented by the epitopes of antibodies AT270, AT8, AT100, AT180, 12E8, and PHF1, and together they could represent a substantial fraction of Tau phosphorylation (∼30% or more; Fig. 5 and (Mair et al., 2016), Fig. 4b). NMR studies revealed some local structural characteristics within this phosphorylated cluster region in Tau (Lippens et al., 2016).

**Fig. 5:**
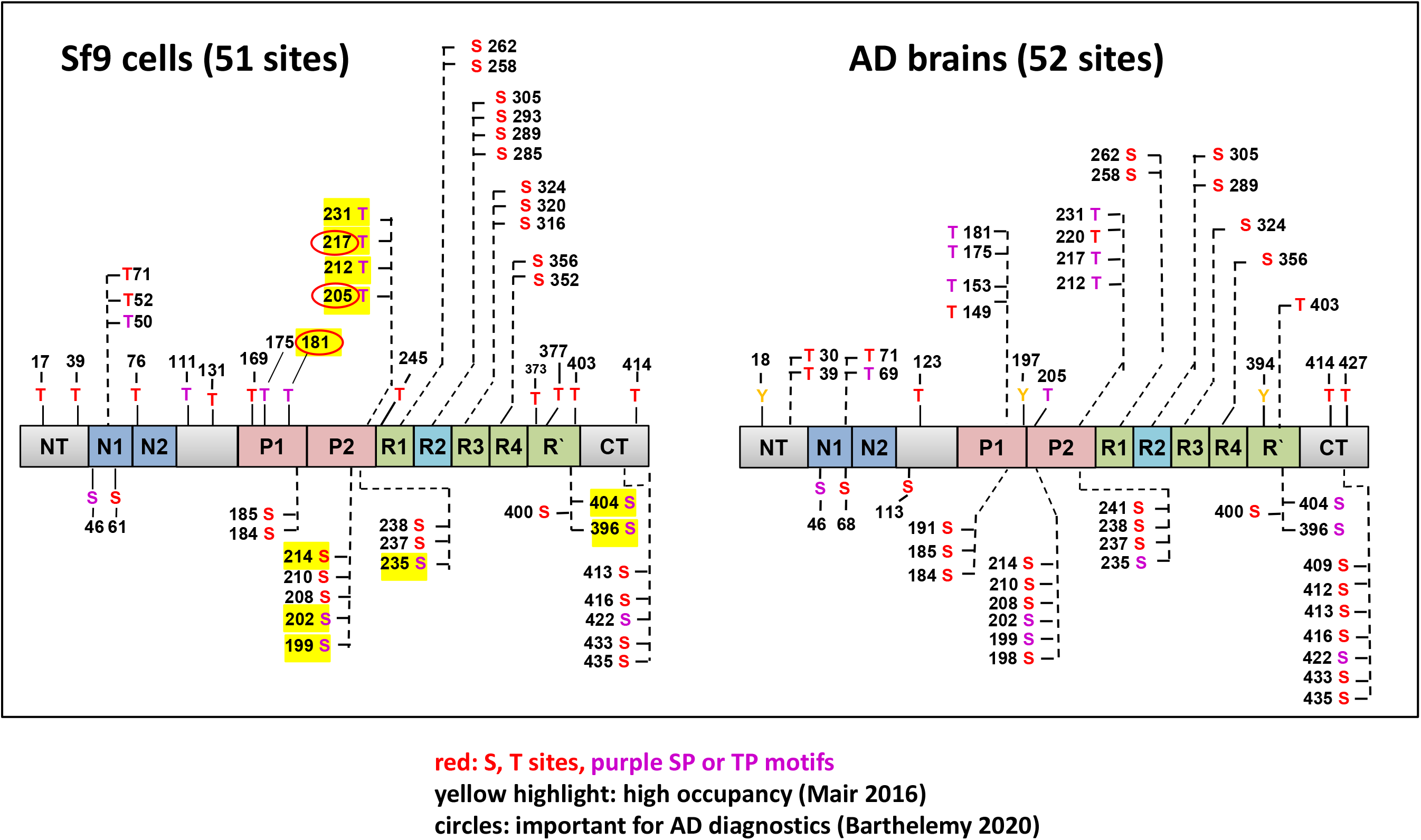
hTau phosphorylation sites observed in Sf9 cells vs. AD brain >50 sites, mainly in C-terminal half, overlap 68%. Bar diagram of full-length human Tau protein with complementary presentation of total phosphorylation sites acquired by Tau in Sf9 cells (left) and phosphorylated sites identified in AD Tau (right).

It is remarkable that all the clusters of phosphorylatable residues mentioned above are located in the C-terminal half of Tau which is responsible for MT interactions and PHF aggregation (Fig. 4). This reflects the asymmetric sensitivity of the domains towards phosphorylation. We can divide up the sequence into an N-terminal 1/3 (approx. residues 1-150, acidic character) and the C-terminal 2/3 (residues 151-441, basic character, including the proline-rich and repeat domains). The total number of 51 detected phosphorylated residues (Fig. 4) represents 51/85=60% of all phosphorylatable residues (S, T, Y). However, the N-terminal 1/3 had only 10/27=37% potential sites phosphorylated, whereas in the C-terminal 2/3 the fraction of phosphorylated vs. potential sites was almost twice as large, 41/58=71%. Since all Tau domains are largely unstructured and mobile in solution (Mukrasch et al., 2009), it is unlikely that the difference in sensitivity to kinases arises from a protection of sites via folding (especially since the N-terminal part is known for its role as a “fuzzy coat” (Jakes et al., 1991). Other conceivable explanations are (1) some kinases of Tau may prefer target motifs of basic character, and/or (2) kinases of Tau may first bind to acidic domains before phosphorylating other parts of the Tau chain. Direct evidence for this scenario comes from NMR studies of the kinase MARK which binds to the N-terminal region of Tau and then phosphorylates the KXGS motifs in the repeat domain (Schwalbe et al., 2013).

### Tau phosphorylation versus aggregation in AD

The hotspots of Tau phosphorylation in Sf9 cells coincide well with the epitopes of the antibodies mentioned above, raised initially against PHFs purified from Alzheimer brain. The coincidence of aggregated and phosphorylated states in AD tissue led to the hypothesis that (hyper-) phosphorylation predisposes Tau for aggregation, and set off an extended search for kinases of Tau (Martin et al., 2013, Kosik and Shimura, 2005). However, this effort remained inconclusive, as no kinase with pronounced pro-aggregant activity was found (in contrast to FTDP17 Tau mutations which are clearly pro-aggregant), and some phosphorylation sites were even protective against aggregation (Schneider et al., 1999). The high solubility of Tau from Sf9 cells illustrates the lack of a direct causal link between phosphorylation and aggregation. In support of this, although AD-diagnostic antibodies react with Tau monomers from Sf9 cells (see (Tepper et al., 2014), this is not per se an indicator of cell degeneration except in diseases such as AD and transgenic mice with Tau pathology. Indirect causal links may include phosphorylation-dependent conformational changes combined with proteolytic cleavage which exposes aggregation-prone domains of Tau (e.g. repeat domain). Examples are cleavage by calpain or caspases (Wang et al., 2007, Flores et al., 2018) or non-enzymatic cleavage in aging cells which causes the heterogeneous “smear” in SDS gels of AD Tau (Watanabe et al., 2004). In support of this, incipient neurodegeneration correlates with the appearance of soluble tau oligomers rather than Tau-containing tangles (Lasagna-Reeves et al., 2012, Kaniyappan et al., 2017).

### Tau phosphorylation in cell models

As mentioned earlier, the functions of Tau in cytoskeletal remodeling, studied initially in PC12 cells and neurons (Drubin et al., 1985), were well replicated in other eukaryotic cell types of human or animal origin including Sf9 cells (Knops et al., 1991). However, one may ask whether this also involves similar P-sites. In previous work we addressed this issue by studying CHO cells transfected with human Tau (2N4R), and a mouse neuroblastoma cell line (LAN-5) expressing endogenous Tau (Illenberger et al., 1998). The cells were metabolically labeled with ^32^P, followed by phosphopeptide mapping. There was a striking overlap of P-sites with the current results, especially involving the clusters of P-sites listed above. Phosphorylation became enhanced during mitosis, corresponding to the activation of the cell cycle kinase cdc2, a proline-directed kinase. Note that this phosphorylation occurred during a physiological process, without any sign of pathology or Tau aggregation, indicating that the pattern of phosphorylation serves a normal biological role. The same conclusion can be drawn from the analysis of hibernating animals where retraction and regrowth of neuronal networks are accompanied by similar changes in Tau phosphorylation (von der Ohe et al., 2007, Arendt et al., 2003).

### Other types of PTMs in Sf9 cells

So far we have focused on phosphorylation because it is the most conspicuous feature of Tau modification and is at the focus of biomedical research. On the other hand, research on Tau has unearthed a large variety of modifications, including chemical modifications of residues, cleavage by various proteases, or specific interactions with other regulatory components (Morris et al., 2015). For the case of Sf9 cells, the striking observation is the very low level of other modifications that have received attention in the field:

**Phosphorylation of Tyr** is absent or below detectability, in contrast to the abundance of Ser/Thr phosphorylation. The 5 Tyr-residues in Tau (residues 18, 29, 197, 310, 394) can be phosphorylated by common Tyr kinases in the brain, e.g. Fyn, Src, Lck, Abl, and the proline-rich regions contain several binding sites for associated SH3 domains of the type PXXP (Derkinderen et al., 2005, Hanger et al., 2009, Ittner et al., 2010). This makes Tau interesting as a carrier of cellular signaling. However, at least in Sf9 cells, the Tyr-dependent signaling affects at best a minute fraction of Tau.

**Acetylation of Lys:** This modification is important for gene regulation, but occurs in the cytoplasm as well, notably on microtubules and associated proteins (e.g. K174 in Tau) and has been implicated in neuronal damage of aging cells (Sohn et al., 2016). The low level of acetylated Tau can be explained by the high activity of HDACs in Sf9 cells which makes them highly radiation-resistant (Sharma et al., 2016).

**Ubiquitination of Lys** is a frequent Tau modification in aging or AD brains and reflects the cell’s attempt to remove misfolded or aggregated Tau (Kosik and Shimura, 2005, Yu et al., 2019). Ubiquitin binding sites have recently been revealed by high-resolution cryo-EM imaging of Tau filaments (Arakhamia et al., 2020). However, this modification is not detectable in Sf9 cells. A simplified interpretation of these results is that Sf9 cells are “healthy” in a cellular sense: Ubiquitination of Tau is very low, consistent with the absence of aggregates. Likewise, acetylation is low as well. The phosphorylation tone is high, comparable with that of fetal neurons in absence of aggregates. Further evidence for the healthy state is that the Sf9 cells develop extensions analogous to those of differentiating neurons, stabilized by bundles of parallel microtubules and microtubule associated Tau (Biernat and Mandelkow, 1999, Knops et al., 1991). This implies that the increased Tau phosphorylation is not necessarily detrimental and occurs under both physiological and pathological conditions.

### Tau-associated proteins

In both cell types studied here (*E*.*coli* and Sf9), LC-MS/MS and native MS revealed traces of numerous proteins which remained associated with Tau through the purification (see Tables 3 and 4, Figures 2 and 3). This agrees well with the multiple published interactions of Tau (Mandelkow and Mandelkow, 2012). Many of the accessory proteins belong to the class of ribonucleoprotein complexes (Gunawardana et al., 2015, Maziuk et al., 2017). Examples are proteins involved in mRNA metabolic processes, e.g. translation initiation, and translation elongation (Wheeler et al., 2019). Tau expression can differentially shift both the transcriptome and the nascent proteome, and the synthesis of ribosomal proteins is reversibly dependent on Tau levels (Meier et al., 2016, Koren et al., 2019). The presence of DNA binding proteins is reminiscent of nuclear functions of Tau in DNA protection under stress conditions (Multhaup et al., 2015). Finally, there are numerous examples of interactions between Tau and chaperones (Karagoz et al., 2014), a class that is prominent among the identified associated proteins. For example, the periplasmic chaperone protein Skp from *E*.*coli* interacts with membrane proteins, thus maintaining the solubility of early folding intermediates during passage through the periplasm (Chen and Henning, 1996). Skp is similar to Prefoldin/GimC, a cytosolic chaperone present in eukarya and archaea (Cowan and Lewis, 2001). Prefoldin, a protein chaperone used in protein folding complexes, works as a transfer protein in conjunction with a molecule of chaperonin to correctly fold other nascent proteins, such as the cytoskeletal proteins actin and tubulin.

In summary, by combining the results from native MS analysis of intact Tau and peptide identification using LC-MS/MS we have been able to specify the state of modified Tau in eukaryotic Sf9 cells in comparison to the nearly unmodified state of Tau expressed in *E*.*coli*. Among the identified PTMs on Tau from Sf9 cells, extensive phosphorylation at Ser/Thr residues is by far the dominant modification (up to 20 P_i_ per molecule), whereas other well-known modifications are present only in minute proportions (acetylation, Tyr phosphorylation, ubiquitination). The S/T phosphorylation has a remarkably asymmetric distribution, whereby a low proportion of residues is modified in the acidic N-terminal domain of Tau, in contrast to a high proportion in the basic middle to C-terminal domains, including all of the sites detected by antibodies against Alzheimer phospho-Tau. This indicates that a substantial degree of phosphorylation (achieved by multiple kinases) is presumably a property of normal soluble Tau in eukaryotic cells, reflecting a phosphorylation tonus dominated by net activity of kinases over phosphatases. By implication, this would be expected to affect many other proteins as well, notably natively unfolded proteins accessible to diverse kinases (Oka et al., 2011, Wisniewski et al., 2010). The results may help to identify further markers of disease states in neurons or elsewhere.

## Materials and Methods

### Protein preparation and purification

Tau protein (clone htau40, largest isoform in human CNS, 441 residues, Uniprot Id P10636 isoform F) expressed in *E. coli* or Sf9 cells, prepared and purified as described in (Tepper et al., 2014). Expression in either system yielded Tau concentrations up to ∼230 µM without causing aggregation. In the case of Sf9 cells the expressed Tau protein was purified from cell extracts by making use of the heat stability of Tau. Sf9 Cells were incubated for 3 days at 27 °C and collected directly for preparation of phosphorylated hTau40 protein in lysis buffer (50 mM Tris-HCl, pH 7.4, 500 mM NaCl, 10% glycerol, 1% Nonidet P-40, 5mM DTT, 10mM EGTA, 20mM NaF, 1 mM orthovanadate, 5 µM microcystin, 10 µg/ml each of protease inhibitors leupeptin, aprotinin, and pepstatin) in a ratio of 1 g of Sf9 pellet to 10 ml of lysis buffer. This procedure yielded “P_m_-Tau” (alias P12-Tau in Tepper, 2014; or P-Tau in Mair, 2016). To increase the phosphorylation even further, Sf9 cells were treated before harvesting for 1 h with 0.2 µM okadaic acid (OA; a phosphatase inhibitor, Enzo-LifeScience). Next, after centrifugation the cells were resuspended in lysis buffer and boiled in a water bath at 100°C for 10 min. By this treatment nearly all proteins were denatured and precipitated, except for Tau which stays soluble. The cell debris was removed by centrifugation. The supernatant containing soluble Tau protein was concentrated in Millipore Amicon Ultra-4-centrifugal filter units (molecular mass cutoff of 3 kDa). This procedure yielded “P_h_-Tau” (alias P20-Tau in (Tepper et al., 2014), or PP-Tau in (Mair et al., 2016)). The material was then applied to a size exclusion column Superdex G200 (GE Healthcare) and eluted with PBS buffer (pH 7.4; 1 mM DTT). For further experiments, the fractions containing Tau protein were pooled and concentrated again in Amicon Ultra-4-centrifugal filter units. Finally, the concentrated protein was rebuffered into 200 mM ammonium acetate, pH 7,6.

### Analysis of human Tau proteins by native MS

Purified proteins in 200 mM ammonium acetate, pH 7.6 were analyzed by nanoflow electrospray ionization mass spectrometry using a Synapt HDMS (Waters and MS Vision) equipped with a 32,000 *m/z* range quadrupole. 3 µl of sample were introduced with an in-house manufactured gold-coated capillary needle (borosilicate thin wall with filament, OD 1.0 mm, ID 0.78 mm; Harvard Apparatus). The capillary voltage was set to 1.3 kV and the cone voltage to 180 V. The trap cell was filled with argon gas at a flow rate of 1 ml/min and collisional activation was performed by applying acceleration voltages to the trap cell. The transfer cell was kept at a pressure of about 20% of that in the trap cell and a low acceleration voltage of 5 V. Spectra were calibrated externally using caesium iodide. Charge state spectra were deconvoluted by the program UniDec version 2.7.3 (Marty et al., 2015).

### Identification of peptide sequences and posttranslational modifications by LC-MS/MS

In-solution tryptic digestion of proteins was performed as described previously (Morgenstern et al., 2017). In brief, Tau protein samples were subjected to reduction of cysteine residues with 5 mM Tris(2-carboxyethyl)phosphine, alkylation of free thiol groups with 50 mM iodoacetamide/50 mM NH_4_HCO_3_ and digested with trypsin overnight at 37°C. Peptide mixtures were analyzed by nano-HPLC-ESI-MS/MS using a Q Exactive instrument directly coupled to an UltiMate 3000 RSLCnano HPLC system (both Thermo Fisher Scientific, Dreieich, Germany) equipped with a Nanospray Flex ion source with DirectJunction (Thermo Fisher Scientific). Peptides were washed and preconcentrated on PepMap™ C18 precolumns (5 mm x 300 μm inner diameter; Thermo Scientific) and separated using AcclaimPepMap™ RSLC columns (50 cm x 75 μm inner diameter; pore size, 100 Å; particle size, 2 μm) at a flow rate of 250 nl/min and 40 - 43°C. For peptide elution, binary solvent systems were used consisting of 0.1% (v/v) formic acid (FA) (solvent A) and 0.1% (v/v) FA/86% (v/v) acetonitrile (solvent B) with a gradient of 4-42% solvent B in 50 min, 42 - 95% in 5 min and 5 min at 95%. The Q Exactive was operated with the following parameters: MS survey scans ranging from *m/z* 375 - 1,700 at a resolution (R) of 70,000 (at *m/z* 200), an automatic gain control (AGC) of 3 × 10^6^ ions, and a maximum injection time (IT) of 60 ms. A TOP12 method was used for higher energy collisional dissociation (HCD) of precursor peptides (*z* ≥ 2) in the orbitrap applying a normalized collision energy of 28%, an AGC of 1 × 10^5^ ions and max. IT of 120 ms. The dynamic exclusion time for previously fragmented precursors was set to 45 s.

Data were analysed using Maxquant software version 1.5.5.1 (Tyanova et al., 2016) searching against the sequence of human Tau-F (Uniprot Id P10636-8) and UniProt organism specific sequence databases for *E. coli* and *Spodoptera frugiperda* (version 2018_08, 4,344 and 26,434 entries, respectively) allowing a maximum of four missed cleavages of trypsin, a mass tolerance of 4.5 ppm for precursor and 20 ppm for fragment ions. Methionine oxidation, acetylation of lysine and protein N-terminus as well as phosphorylation at serine, threonine and tyrosine residues were specified as variable modifications and carbamidomethylation of cysteine as fixed modification. Additional searches were performed with an extended range of variable modifications including ubiquitination (diglycine at lysine) and O-GlcNAc modification of serine and threonine. However, these modifications were observed only sporadically and in low intensity and borderline significance and were therefore excluded from the final search. The lists of both peptide and protein identifications were filtered applying a threshold for the false discovery rate of <0.01.

## Acknowledgements

We are grateful to Sabrina Hübschmann (DZNE Bonn) and Bettina Knapp (Univ. Freiburg) for their expert technical assistance, and to Drs. Katharina Tepper DZNE Bonn), Judith Steen (Harvard Med. School, Boston), and Thomas Timm (Univ. Giessen) for critical discussions.

## Funding and additional information

The project was supported by DZNE, MPG, and Katharina-Hardt-Foundation (to EM&EMM) and by the Deutsche Forschungsgemeinschaft (DFG, German Research Foundation) grants FOR1905 and Project-ID 403222702/SFB 1381, as well as the Excellence Strategy (CIBSS – EXC 2189 – Project-ID 390939984) and the Excellence Initiative of the German Federal & State Governments (EXC 294, BIOSS). MS raw data and result files have been deposited to the ProteomeXchange Consortium via the PRIDE repository (Perez-Riverol et al., 2019) and are publicly accessible from its website (http://www.ebi.ac.uk/pride) with the dataset identifier PXD020985.

## Abbreviations and definitions

htau40: Tau-2N4R largest Tau isoform in human CNS, 441 residues
Pi: Phosphate group (on Ser, Thr, or Tyr)
MS: mass spectrometry
LC-MS/MS: Liquid chromatography-mass spectrometry
MALDI TOF-MS: Matrix-assisted laser desorption/ionization time-of-flight mass spectrometry
Sf9 cells: *Spodoptera frugiperda* cells
CSF: Cerebrospinal fluid
HDAC: Histone deacetylase
PTM: posttranslational modification
site occupancy: number of modifications (e.g. phosphorylation) at a given residue position, relative to number of molecules (max. 100%)
overall occupancy: total number of modifications of a certain type (e.g. phosphorylation) per tau molecule (up to ∼20)

---

**Supplement Table S1.**
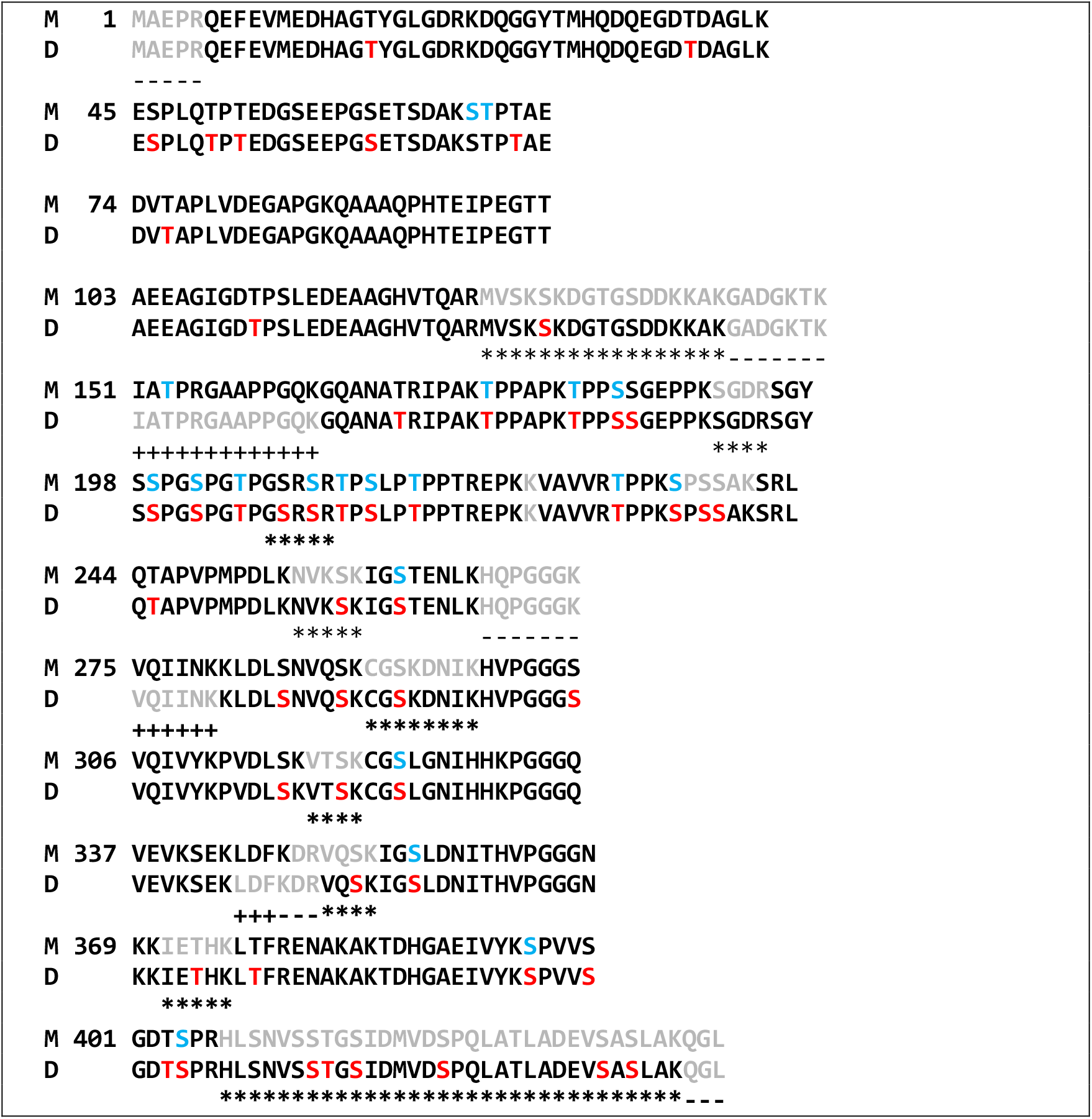
Sequence coverage of Tau peptides in Mair 2016 (M) and current work (D) with all P-sites found. Red letters= P-sites observed in current work. Blue=sites observed in Mair 2016 study. Note that the newly covered sequences in this study (symbol ***) contain 14 observed phosphorylation sites (red letters S, T). Sequences covered previously (Mair2016) but not in current study (symbol +++) contain only 1 observed phosphorylation site (blue, T153). Thus, the excess of new sequence coverage is sufficient to explain the higher occupancy in the current study. grey: missing residues in sequence coverage. *** Gaps in (M) +++ Gaps in (D); --- Gaps in both.

